# Revisiting Sodium Phosphotungstate and Ammonium Molybdate as non-radioactive negative staining agents for single particle analysis

**DOI:** 10.1101/2024.07.19.604222

**Authors:** Monika Gunkel, Arthur Macha, Elmar Behrmann

## Abstract

This study reports the successful replacement of uranyl-based stains by either sodium phosphotungstate or ammonium molybdate for negative staining electron microscopy. Using apoferritin as a test specimen, it is demonstrated that in combination with a facile on-grid fixation step both stains yield comparable images to uranyl formate. Subsequently, using β-galactosidase, it is shown that both stains can also successfully be employed for single particle analysis, yielding virtually indistinguishable results from uranyl formate. As both replacement stains are non-radioactive, they are not subjected to the same handling restrictions as uranyl-based stains. Therefore they are not only cheaper to use, but also make decentralized sample grid preparation – thus directly after purification – accessible to a broader range of scientists.

## 1. Introduction

In recent years cryo-electron microscopy (cryo-EM) has reached a level of maturity allowing it to routinely complement, and sometimes surpass, the capabilities of X-ray crystallography and NMR to uncover the structural basis of protein activity. As with other structural biology tools, the success of cryo-EM relies on successful sample preparation. Often, this becomes the bottleneck of a structural biology project, requiring extensive and time-consuming optimization steps (Takizawa *et al*., 2017; Weissenberger *et al*., 2021). Negative staining is a well-established technique to prepare a wide range of samples for electron microscopy (EM) characterization (Bradley, 1962; Brenner and Horne, 1959; De Carlo and Harris, 2011; Harris, 2015). It is a rapid, robust, and cost-efficient approach, especially compared to cryo-EM. While associated with a range of drawbacks, particularly in terms of achievable resolution and preservation of the native structure, negative stain EM has been demonstrated repeatedly to yield important insights (Fabre *et al*., 2017; Matsuike *et al*., 2023; Pramanick *et al*., 2021; Sasajima *et al*., 2022). For structural biology applications negative stain EM can provide information about general sample quality, regarding particle size distribution, aggregation tendency, homogeneity and abundance of the protein of interest. Moreover, as only neglectable amounts of sample are needed, negative stain EM does not require dedicatedly produced sample material, but can instead be used with left-over material. Therefore it lends itself as a diagnostic tool in cases where the interpretation of cryo-EM datasets has raised doubts about the quality of the input sample. Here, the enhanced image contrast due to embedding into a heavy metal salt film allows to overcome challenges posed by the low particle contrast inherent to cryo-EM. For research questions that do not require high-resolution structural details, such as investigating conditions which allow the formation of protein complexes, determining protein dimensions, and addressing binding sites, negative staining EM has been successfully used as a stand-alone technique (Burgess *et al*., 2004; Fabre *et al*., 2017; Ha *et al*., 2016; Francis *et al*., 2024).

While there is a rich history of chemically diverse staining solutions (Scarff *et al*., 2018), current negative staining EM for single particle analysis (SPA) is almost exclusively relying on either uranium acetate (UA) (Epstein, 2001; Johnson *et al*., 1977) or uranium formate (UF) (Finch, 1964; Leberman, 1965). Both uranium salts yield excellent contrast due to the high Z-number of uranium, and both salts have fine grain sizes allowing to resolve molecular details (Ohi *et al*., 2004). Moreover, the counter-ions of uranium in these salts, acetate and formate, have the advantage of rapidly fixing biological samples during the staining process, with fixation reported to occur in the milliseconds time range (Zhao & Craig, 2003). Still, the widespread use of uranium-based staining solutions not only produces potentially radioactive waste, but has additional drawbacks: firstly, uranium hydroxide will precipitate at neutral and basic pH values, and thus stains are commonly prepared as highly acidic solutions that potentially interfere with the biological sample of interest. Secondly, due to its classification as a radioactive substance obtaining, handling, and disposing of uranium salts can be challenging from a legal and administrative point of view, which often translates into high costs. These administrative burdens also complicate decentralized sample preparation, so that purified protein samples often must be brought to central EM labs before they can be stained, which can be an issue with instable samples. Given these drawbacks, it is not surprising that alternative stains are sought after. However often these studies focus on the staining and preservation of subcellular structures, and not on SPA applications (Benmeradi *et al*., 2015; Inaga *et al*., 2007; Nakakoshi *et al*., 2011; Hosogi *et al*., 2015; Kuipers & Giepmans, 2020; Scarff *et al*., 2018).

Here, we report the successful application of two stains that were used for SPA before the advent of UF and UA, namely sodium phosphotungstate (SPT) (Brenner & Horne, 1959) and ammonium molybdate (AMo) (Bohonek, 1974; Mannella & Frank, 1984). We show that by including a facile on-grid fixation step both SPT and AMo can be used to visualize the 440 kDa capsid assembly of apoferritin. Furthermore, we demonstrate that β-galactosidase, a common test specimen in the field of cryo-EM, can be faithfully reconstructed using both stains.

## 2. Materials and methods

### 2.1. Protein expression and purification

*Mus musculus* apoferritin heavy chain (pET24a-mmFTH1) was expressed in *Escherichia coli* BL21(DE3) competent cells. BL21(DE3) transformed with pET24a-mmFTH1 were cultured in LB medium (10 g/L tryptone, 5 g/L yeast extract, 10 g/L NaCl) supplemented with 50 µg/mL kanamycin. After reaching an OD_600_ of 0.5, IPTG was added to a final concentration of 1 mM to induce expression. Cells were further cultured for 4 h at 37 °C, and then collected by centrifugation at 3000 rpm for 15-20 min at 4 °C. Cell pellets were resuspended in lysis buffer (20 mM HEPES-NaOH pH 7.5, 300 mM NaCl, 1 mM MgSO_4_) supplemented with 100 mg/mL lysozyme, followed by sonication (Power 200 W, amplitude 60 %, C 100 %, Sonotrode diameter 7 mm, 4 s work / 8 s break, 30 cycles) on ice. After clarifying by centrifugation at 70000 *g* for 30 min at 4 °C, the supernatant was heated for 10 min to 70 °C, then centrifuged at 70000 *g* for 30 min at 4 °C. 8.4 g Ammonium sulfate was added to the second supernatant to a final concentration of 52.5 %. The solution was centrifuged at 50000 *g* for 20 min at 4 °C. Afterwards the supernatant was discarded, and the pellet resuspended in a final volume of 2 mL sample buffer (20 mM HEPES-NaOH, pH 7.5, 300 mM NaCl). The sample was further purified by size exclusion chromatography using a HiLoad 16/600 Superdex 200 pg gel filtration column using sample buffer. Fractions containing apoferritin were snap frozen in liquid nitrogen and stored at -80 °C until further use.

*E. coli* ß-galactosidase was purchased from Sigma Aldrich (catalog number G5635-3KU) as lyophilized protein. 2.5 g of the lyophilized protein were dissolved in 250 µL sample buffer (25 mM Tris/HCl pH 8.0, 50 mM NaCl, 0.5 mM TCEP) and aggregates were removed by centrifugation. The sample was further purified by size exclusion chromatography using a Superdex 200 increase 10/300 GL filtration column using sample buffer. Fractions containing ß-galactosidase were snap frozen in liquid nitrogen and stored at -80 °C until further use.

### 2.2. Preparation and storage of staining solutions

UF was prepared by dissolving 2 % (w/v) UF (Science Services, CAS #6984-95-1) in boiling ddH_2_O. The solution was stirring in darkness for 5 min. Then 6.4 µL 4M NaOH per 1 mL UF solution were added and stirred for another 5 min in darkness. The solution was filtered through a 0.22 µM filter, and 200 µL aliquots were shock frozen in liquid nitrogen and stored at -80 °C.

AMo was prepared by dissolving 1 % (w/v) AMo (Sigma Aldrich, CAS #13106-76-8) in ddH_2_O. The pH was set with ammonium hydroxide to pH 7.0. 200 µL aliquots were shock frozen in liquid nitrogen and stored at -80 °C.

SPT was prepared by dissolving 2 % (w/v) SPT (Sigma Aldrich, CAS #312696-30-3) in ddH_2_O. The pH was set with sodium hydroxide to pH 7.0. 200 µL aliquots were shock frozen in liquid nitrogen and stored at -80 °C.

For storage recommendations see the Supplementary Information and Figure S1.

### 2.3. Negative staining with UF

Continuous carbon grids (Quantifoil, Cu 200 mesh) were subjected to glow discharge using a Zepto Plasma Cleaner (Diener) for 30 seconds to clean the surface and render it hydrophilic. 3 µL of protein solution, at a concentration of 0.4 mg/mL for apoferritin and 2.1 mg/mL for β-galactosidase, were incubated on the grid for approximately 1 min at room temperature, then excess protein solution was blotted away using filter paper, taking care to leave a thin liquid film on the grid. Using self-locking forceps (Dumoxel, N5) the grid was then washed by dipping it onto a droplet of protein buffer, and blotting away the buffer using filter paper, taking care to leave a thin liquid film. This step was repeated three times. Next, the grid was briefly dipped onto a droplet of staining solution, followed by immediate blotting of excess liquid with filter paper. The grid was then placed onto fresh droplets of staining solution and incubated for 1 min. Excess stain was carefully removed with filter paper, taking care to leave a thin liquid film on the grid. The remaining stain solution was then rapidly dried by using a gentle, indirect stream of a hair dryer without heating. Grids were stored in the dark in a dry atmosphere until further usage.

### 2.4. Negative staining with AMo or SPT

Continuous carbon grids (Quantifoil, Cu 200 mesh) were subjected to glow discharge using a Zepto Plasma Cleaner (Diener) for 30 seconds to clean the surface and render it hydrophilic. 3 µL of protein solution, at a concentration of 0.4 mg/mL for apoferritin and 2.1 mg/mL for β-galactosidase, were incubated on the grid for approximately 1 min at room temperature, then excess protein solution was blotted away using filter paper, taking care to leave a thin liquid film on the grid. Using self-locking forceps (Dumoxel, N5) the grid was then washed by dipping it onto a droplet of protein buffer, and blotting away the buffer using filter paper, taking care to leave a thin liquid film. This step was repeated 3 times. Next, the grid was briefly dipped onto a droplet of fixation solution 0.15% (w/v) glutaraldehyde in ddH_2_O, Sigma Aldrich, CAS #111-30-8) followed by immediate blotting of excess liquid with filter paper. The grid was then placed onto a fresh droplet of fixation solution and incubated for 5 min. Excess fixation solution was carefully removed with filter paper. Next, the grid was briefly dipped onto a droplet of staining solution, followed by immediate blotting of excess liquid with filter paper. The grid was then placed onto fresh droplets of staining solution and incubated for 1 min. Excess stain was carefully removed with filter paper, taking care to leave a thin liquid film on the grid. The remaining stain solution was then rapidly dried by using a gentle, indirect stream of a hair dryer without heating. Grids were stored in the dark in a dry atmosphere until further usage.

For a detailed, step-by-step guide see the Supplementary Information and Figure S2.

### 2.5. EM data collection

EM data was acquired using a Talos L120C (Thermo Fisher Scientific) electron microscope equipped with a LaB_6_ emitter operated at 120 kV. Images were collected automatically using EPU (Thermo Fisher Scientific, version: 2.12.1.2782REL) on Ceta16M CMOS detector with a calibrated pixel size 1.86 Å/px. Defocus values were set to range from -0.3 to -2.0 μm.

### 2.6. Image processing, particle quantification, and data analysis

Image processing was performed using cryoSPARC 4.4 (Punjani *et al*., 2017). Random subsets of 25 micrographs were used for each staining condition to determine the optimal value for the amplitude contrast by comparing experimental to simulated CTF oscillations. Using this identified value (UF: 0.35, AMo: 0.15, SPT: 0.10) defocus and other CTF-related values were then calculated for the complete datasets. Only high-quality micrographs with low astigmatism and good agreement between experimental and calculated CTFs were further processed.

For apoferritin a total of nine grids were prepared during three independent staining sessions, with each session using freshly prepared staining solution. During each session a single grid was prepared for each of the three stains, namely UF, SPT and AMo. From each grid, several micrographs were acquired. Of these, 5 micrographs were chosen at random, resulting in 15 micrographs for UF, SPT and AMo, respectively. On these micrographs particles were picked manually and classified as either “dark core”, “bright core”, or “ambiguous” by hand. Based on this classification, the percentage of either “dark core”, “bright core” or “ambiguous” was calculated for each image independently to account for the different particle counts on each image. Finally, the mean of all particle numbers of a staining solution was calculated together with the standard deviation between the individual images.

For β-galactosidase, each dataset was limited to 163 high-quality micrographs. On these, putative particles were automatically picked based on the expected protein diameter of 180 Å, extracted, and subjected to reference-free 2D classification. Representative 2D classes were then used for a template-based picking approach, particles extracted again, subjected to reference-free 2D classification to exclude artefacts, and subsequent 3D classification using C1 symmetry to identify high-quality particles. The particle population yielding a 3D volume showing four defined subunits was further refined using the homogeneous refinement strategy with enforcing D2 symmetry. To account for the limiting grain size of the negative stain salts and minimize overfitting we limited the alignment resolution for 3D classification to 15 Å, and to 10 Å for the homogeneous refinement. Final maps were low-pass filtered to 10 Å for comparability. The atomic model of *E. coli* β-galactosidase (PDB ID 6X1Q) was fitted into the electron density maps using a rigid body strategy as implemented in ChimeraX (version 1.8-rc2024.06.06) (Meng *et al*., 2023).

For further details see Figures S3, S4 and S5.

## 3. Results

### 3.1. SPT and AMo require on-grid fixation to faithfully stain apoferritin

As a first step to evaluate whether SPT and AMo are suitable replacements for the established uranyl-based stains, we investigated their ability to fixate and stain the iron-storage protein apoferritin (Hamaguchi *et al*., 2019; Kayama *et al*., 2021; Wu *et al*., 2020). Ferritin is a spherical capsid, with an outer diameter of 12 nm and an inner cavity with a diameter of 7 nm, comprising 24 subunits in Eukaryotes (Massover, 1993; Narayanan *et al*., 2019). After staining with UF we observed mainly circular objects with a diameter of 12 nm that have an electron dense, dark core with a diameter of approximately 7 nm (particles with dark core: 98 ± 2 %) (Figure 1 (*a*) and Figure S6 (*a*)).

**Figure 1.**
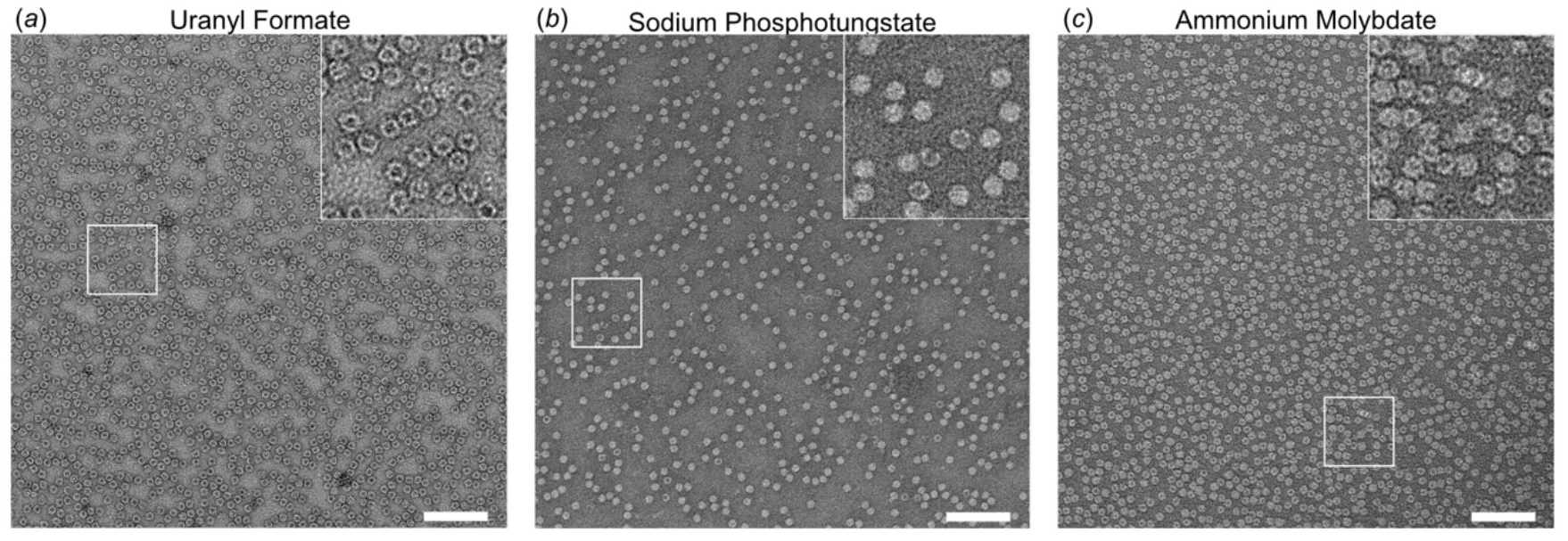
UF stained apoferritin particles feature an electron-dense core, while SPT or AMo stained particles do not. Representative raw micrographs of apoferritin stained with (*a*) UF, (*b*) SPT, (*c*) or AMo. Scale bars are 100 nm. The insets show the boxed area at 2.5 x magnification. For micrographs of SPT and AMo staining without prior on-grid fixation see Figure S7. For a quantification of particles according to their core area see Figure S6.

Conversely, if using the same staining protocol but replacing UF with either SPT or AMo we barely observed defined apoferritin capsids as particles appear fuzzy (Figure S7). Assuming a disassembly of the capsids during the staining procedure, we included an on-grid fixation step prior to the staining step into our staining protocol (Figure 2, Supplementary Information and Figure S2).

**Figure 2.**
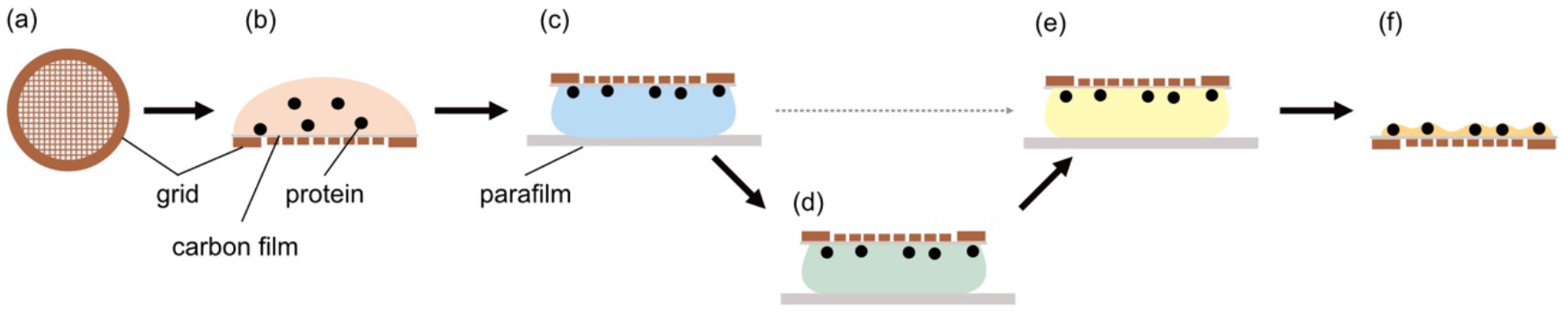
Brief overview of the negative staining workflow. (*a*) Starting from glow discharged sample carrier grids with a continuous support film, (*b*) protein in buffer solution is incubated on the grid surface. (*c*) Grids are then washed with sample buffer. (*d*) For AMo and SPT negative staining an on-grid fixation step using 0.15% glutaraldehyde solution is employed to fix the sample, (*e*) before staining using a droplet of staining solution. (*f*) Afterwards the stain is rapidly dried to create an amorphous film around the sample. For a detailed step-by-step guide see Figure S2.

This additional fixation step resulted in a marked improvement of the particle quality, revealing circular objects with a diameter of 12 nm that. While 98 ± 2 % of UF stained particles have a dark core, most SPT and AMo stained particles do not show a dark core (SPT staining, particles with dark core: 4.3 ± 6.4 %; AMo staining, particles with dark core: 9.4 ± 5.1 %) (Figure 1 (*b, c*), Figure S6 (*b, c*)).

### 3.2. SPT and AMo can be used to assess sample quality of β-galactosidase

As a proof-of-concept that SPT and AMo are also suitable for SPA applications, we relied on the well-characterized protein β-galactosidase which is a 464 kDa homotetramer with D2 symmetry (Jacobson *et al*., 1994) and is commonly used both in biochemical assays and as a resolution standard for cryo-EM. Individual β-galactosidase molecules could be readily distinguished from the background for all three tested stains, although particles did appear slightly fainter for SPT and AMo staining compared to the established UF stain (Figure 3).

**Figure 3.**
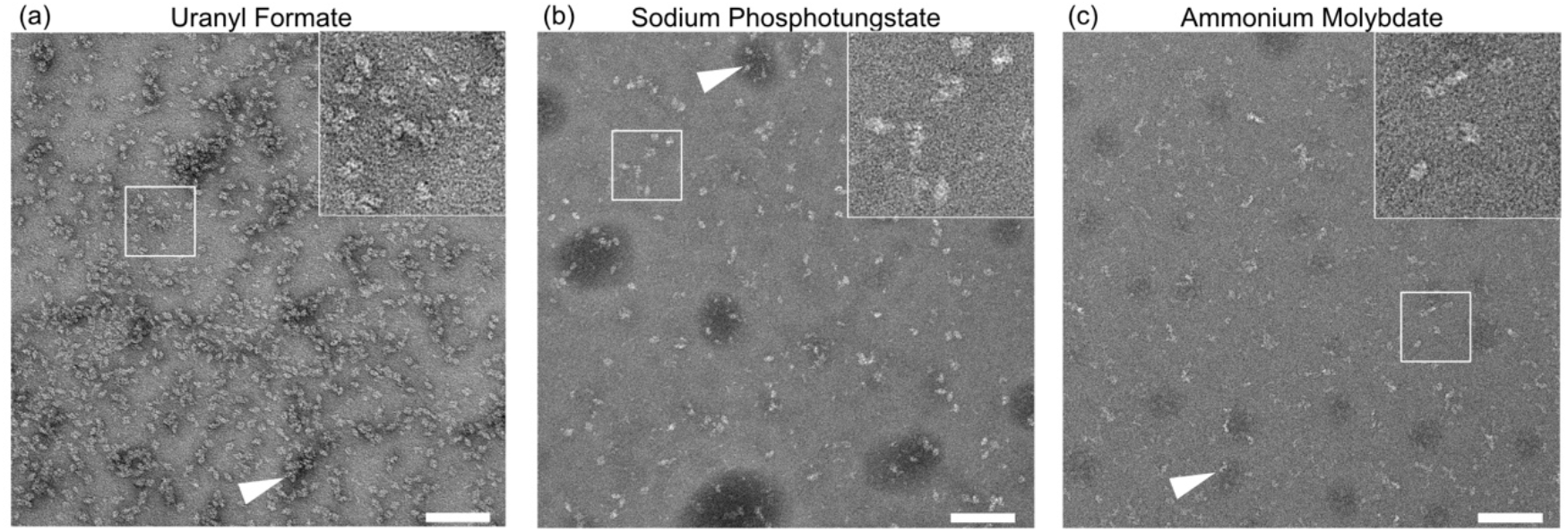
SPT or AMo staining allows assessing the quality of β-galactosidase preparations similar to UF stain. Representative raw micrographs of β-galactosidase stained with (*a*) UF, (*b*) SPT, (*c*) or AMo. Scale bars are 100 nm. The insets show the boxed area at 2.5 x magnification. Instances of stain accumulation, commonly associated with protein aggregates, are marked by arrows.

Nevertheless, all three stains allowed for a rapid assessment of the sample quality. On the micrographs β-galactosidase tetramers could be visually differentiated from smaller objects, such as incomplete assemblies or contaminants and from stain-dense clusters of particles that are commonly associated with (micro-)aggregates (Chari *et al*., 2015) (Figure 3, white arrows). We noted that the observed particle density is lower for SPT and AMo compared to UF, although identical protein concentrations and absorption times were used.

### 3.3. SPT and AMo can be used to reconstruct the 3D structure of β-galactosidase

We next processed data from all three staining conditions to evaluate the capability of SPT and AMo, compared to UF, to preserve the native structure of β-galactosidase (Figures S3, S4, S5). In line with our observation of the raw micrographs, we note that CTF correction for both SPT and AMo worked best assuming a lower amplitude contrast as for UF (in our hands: 0.10 for SPT, 0.15 for AMo, 0.35 for UF). Despite this difference, all three staining conditions yielded convincing 2D class averages that clearly showed the boundaries of the individual subunits. As expected for negative staining, secondary structure elements were not visible in these 2D class averages. Ultimately, all three stains yielded comparable 3D density maps at resolutions around 10 Å that allowed for rigid-body fitting of the known molecular model of the β-galactosidase tetramer (Merk *et al*., 2020) (Figure 4).

**Figure 4.**
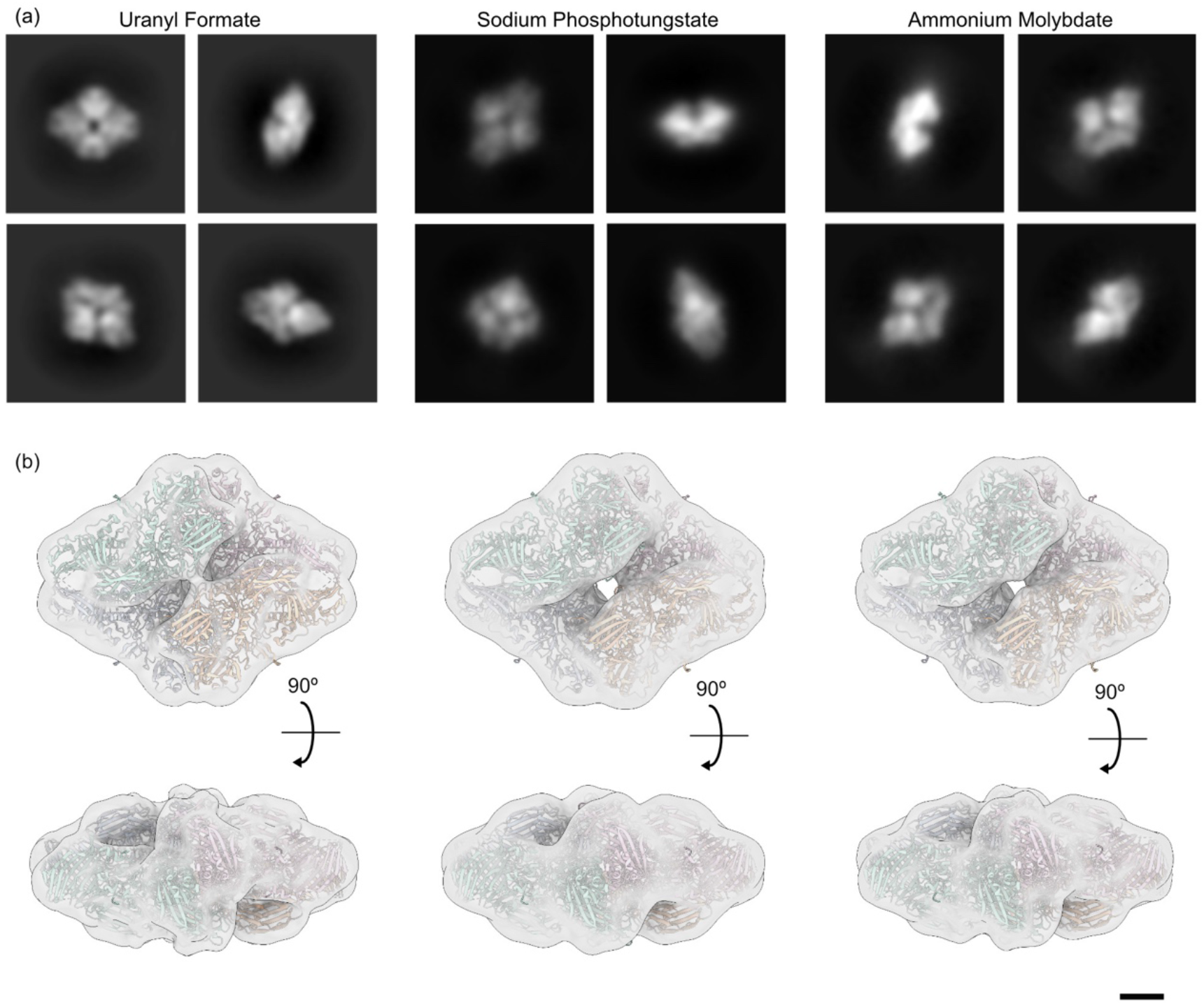
UF, SPT and AMo all allow obtaining faithful 3D reconstructions of β-galactosidase by single particle analysis. (*a*) Selected 2D class averages β-galactosidase obtained from grids prepared by either UF, SPT or AMo negative staining. (*b*) Surface representation of 3D density maps calculated from these datasets, filtered to 10 Å for comparability. The atomic model of β-galactosidase (PDB ID 6X1Q, coloured by individual subunits) was rigid-body fitted to each map. Scale bar is 25 Å. For the complete 3D reconstruction workflow of each condition see Figures S3, S4 and S5.

## Discussion

The majority of structures submitted to the Electron Microscopy Data Bank (EMDB) originates from uranyl-based stains. While this number is potentially skewed by publication bias, since researchers new to the field are more likely to employ well-established and documented stains, there are objective reasons for choosing uranyl-based stains, such as a strong beam scattering effect (Masamichi, 2011), a fine grain size (Haschemeyer & Myers, 1972), and the strong fixating properties of formate and acetate (Zhao & Craig, 2003). However, uranium is a radioactive material, and it requires a low pH to prevent crystallization of the staining solution (Cao et al, 2011). In contrast, SPT and AMo are based on the non-radioactive elements tungsten and molybdenum, which also show adequate solubility in solutions at neutral pH values, with SPT having been reported to be stable in the pH range of 6 to 9 (Scarff *et al*., 2018), and AMo in the pH range of 5 to 7 (Bohonek, 1974). As a proof-of-concept we tested both SPT and AMo stains with two different proteins, apoferritin and β-galactosidase, and compared results to the established UF stain. When working with apoferritin, we observed that the lack of an organic counter-ion with protein fixating properties required us to include a simple on-grid fixation step into our staining procedure (Figure 2, Supplementary Information, Figure S2) in order to stabilize both proteins (Figure 1, 3, S7). The on-grid fixation yielded homogeneous particle distribution on the grids, while using in-solution fixation instead (Harris, 1999) in our hands resulted in aggregation of the test proteins (data not shown). Presumably, the absorption of the protein to the carbon film minimizes the risk of inter-molecular crosslinks.

While apoferritin stained by UF appeared as particles with a dark core, implying that the central cavity of the capsid-like structure was filled with staining solution, the majority of particles stained by either SPT or AMo did not (Figure 1 (*a, b, c*), Figure S6). This difference in staining has been observed before (Harris, 1982). Possibly, the different behavior is due to the pH difference of the staining solutions. It has been reported that low pH solutions can cause opening of the pores of apoferritin, putatively allowing uranium ions to penetrate the capsid (Mollazadeh *et al*., 2022). While SPT and AMo have been reported to have approximately twice the grain size of UF (Harris, 2006; Hosogi *et al*., 2015), we do not observe any resolution-limiting effect compared to UF when working with the tetrameric protein β-galactosidase (Figures S3, S4, S5), and the resulting 3D density maps are virtually identical (Figure 4). Moreover, the apparently lower contrast of SPT and AMo (Figure 1, 3), likely due to the lower atomic numbers of tungsten and molybdenum compared to uranium, did not negatively affect particle picking, 2D classification or 3D reconstruction. The correct amplitude contrast for processing SPT and AMo data was easy to determine by systematically varying this value for a small subset of micrographs during the CTF determination step.

In combination with the easy to integrate on-grid fixation step, SPT and AMo offer adequate contrast and structure preservation, establishing themselves as viable alternatives to uranyl salts for sample preparation for SPA applications. Both SPT and AMo can be utilized at higher pH ranges, permitting the imaging of proteins at neutral pH, in contrast to UF which has limited solubility at non-acidic pH values. Additionally, as non-radioactive alternatives, SPT and AMo can be used in any laboratory without the administrative complexities associated with the purchase and disposal of radioactive materials. This advantage enables researchers to negatively stain their proteins directly after preparation within their own laboratory, eliminating the need to transport samples to an external, possibly remote, EM facility.

## Data availability

The negative stain EM density maps have been deposited in the EMDB under the accession codes EMD-51069, EMD-51071, EMD-51072

## Acknowledgements

We acknowledge access to the cryo-EM infrastructure of StruBiTEM (Cologne, funded by DFG Grant INST 216/949-1 FUGG), and to the computing infrastructure of CHEOPS (Cologne, funded by DFG Grant INST 216/512/1 FUGG). We thank Alexandra Schneider and Jennifer Lange for the preparation of apoferritin and β-galactosidase. We thank Sandra Jordan for contributions at an early stage of the project. We thank Dr. Kristina Barragán Sanz for the kind donation of the pET24a-mmFTH1 plasmid. The authors declare no competing interests.

## Author contributions

M.G. conceived the study; M.G. and A.M prepared samples; M.G. performed electron microscopy; M.G. analyzed data; M.G. performed 3D reconstruction; M.G. and E.B. validated data; M.G. prepared figures; M.G. and E.B. wrote the manuscript with input from all authors.

