## Supplementary Information for "Revisiting Sodium Phosphotungstate and Ammonium Molybdate as non-radioactive negative staining agents for single particle analysis"

### Supporting information

##### S1. Detailed step-by-step workflow for negative staining with SPT or AMo

For a graphical illustration of the workflow see Figure S2.

1. Continuous carbon grid (Quantifoil, Cu 200 mesh) was subjected to glow discharge using a Zepto Plasma Cleaner (Diener) for 30 seconds to clean the surface and render it hydrophilic.

2. All droplets for staining are placed on a clean piece of parafilm: 3 buffer droplets, 2 GA droplets, 2 stain droplets (Figure S2 *f*).

3. The grid was picked up using self-locking forceps (Dumoxel, N5). 3 µl of protein solution, at a concentration of 0.4 mg/ml for apoferritin and 2.1 mg/ml for β-galactosidase, was applied to the grid that is held by the forceps. Protein solution was incubated on the grid for approximately 1 min at room temperature (Figure S2 *a, g*).

4. Excess protein solution was carefully blotted away using filter paper, taking care to leave a thin liquid film on the grid (Figure S2 *k, l*). Critical: In this and all subsequent blotting steps care is taken that there is always a thin film left and the grid never gets fully dry!

5. Still holding the grid in the self-locking forceps, the grid was washed by dipping it onto a droplet of protein buffer and gently moving around on the droplet surface for 2 sec, immediate blotting away using filter paper afterwards. This washing was repeated 3 times using a fresh droplet each time (Figure S2 *b, h*).

6. Next, the grid was briefly dipped onto a droplet of fixation solution (0.15% glutaraldehyde (Sigma Aldrich CAS #111-30-8, dissolved in $ddH_2O$) (Figure S2 *c, h*), followed by immediate blotting of excess liquid with filter paper (Figure S2 *k*).

7. The grid was then placed onto a fresh droplet of fixation solution and incubated for 5 minutes (Figure S2 *c, i*). For this the grid was released from the self-locking forceps and

placed onto the surface of the GA droplet. After the incubation time the grid was picked up from the GA droplet and excess fixation solution was carefully removed with filter paper.

8. Next, the grid was briefly dipped onto a droplet of staining solution (Figure S2 *d, h*), followed by immediate blotting of excess liquid with filter paper.

9. The grid was then placed onto a fresh droplet of staining solution and incubated for 1 min (Figure S2 *d, i*). For this the grid was released from the self-locking forceps and placed onto the surface of the staining droplet. Excess stain was carefully removed with filter paper.

10. The remaining stain solution was then rapidly dried on the grid surface by using a gentle, indirect stream of a hair dryer without heating function (Figure S2 *m*). The gentle steam of the hair dryer blows the remaining staining solution to one side of the grid, so that we get a gradient from thin to thick stain on the grid. This allows for selecting areas of ideal stain thickness, similar to what is done for cryo-EM samples.

##### S2. Guide to storing SPT and AMo staining solutions

To test how stable SPT and AMo are under different storing conditions, stains were prepared, aliquods of 200 µl were used:

- freshly prepared
- after 30 days storage at 4 °C
- after snap freezing in liquid nitrogen, storage at -20 °C for 30 days, thawing
- after snap freezing in liquid nitrogen, storage at -80 °C for 30 days, thawing

As we could not identify obvious differences in the staining behavour (Figure S1) we recommend preparing a larger batch of staining solution and storing 200 µL at either -20 °C or -80 °C until use. Independent of the storage conditions, the staining solutions are stable, show no stain aggregates and comparable staining results (Figure S1).

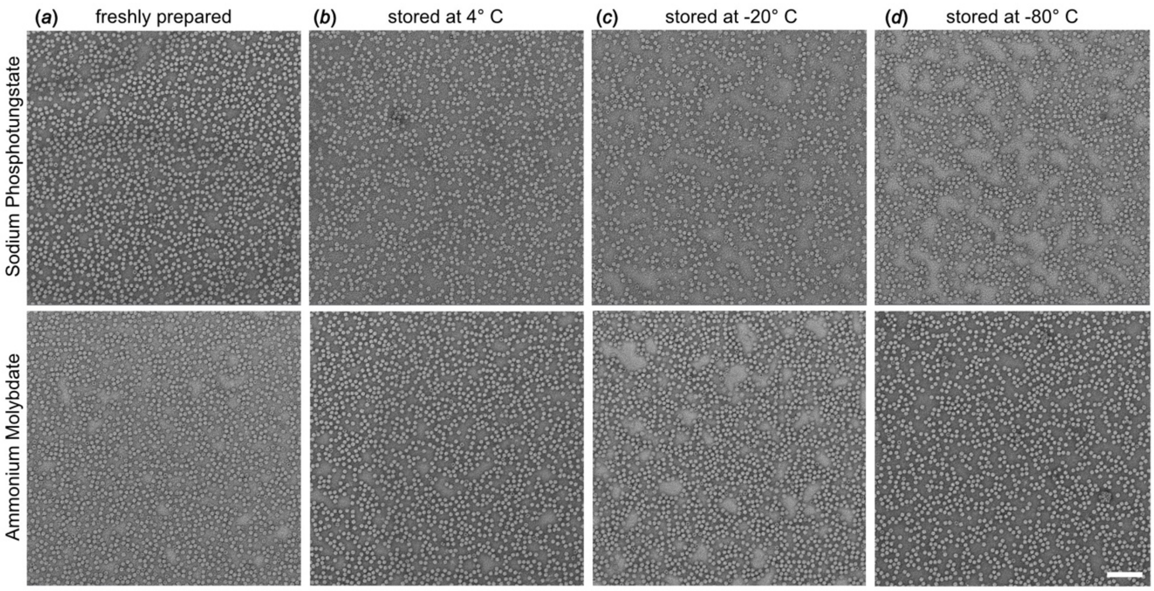


**Figure S1** **SPT and AMo are stable under different storing conditions.** All images show apoferritin stained with either SPT or AMo. Before staining, the staining solutions were (*a*) freshly prepared, (*b*) prepared and stored at 4 °C for 30 days, (*c*) prepared, snap frozen in liquid nitrogen and stored at -20 °C, (*d*) prepared, snap frozen in liquid nitrogen and stored at -80 °C. Scale bar 100 nm.

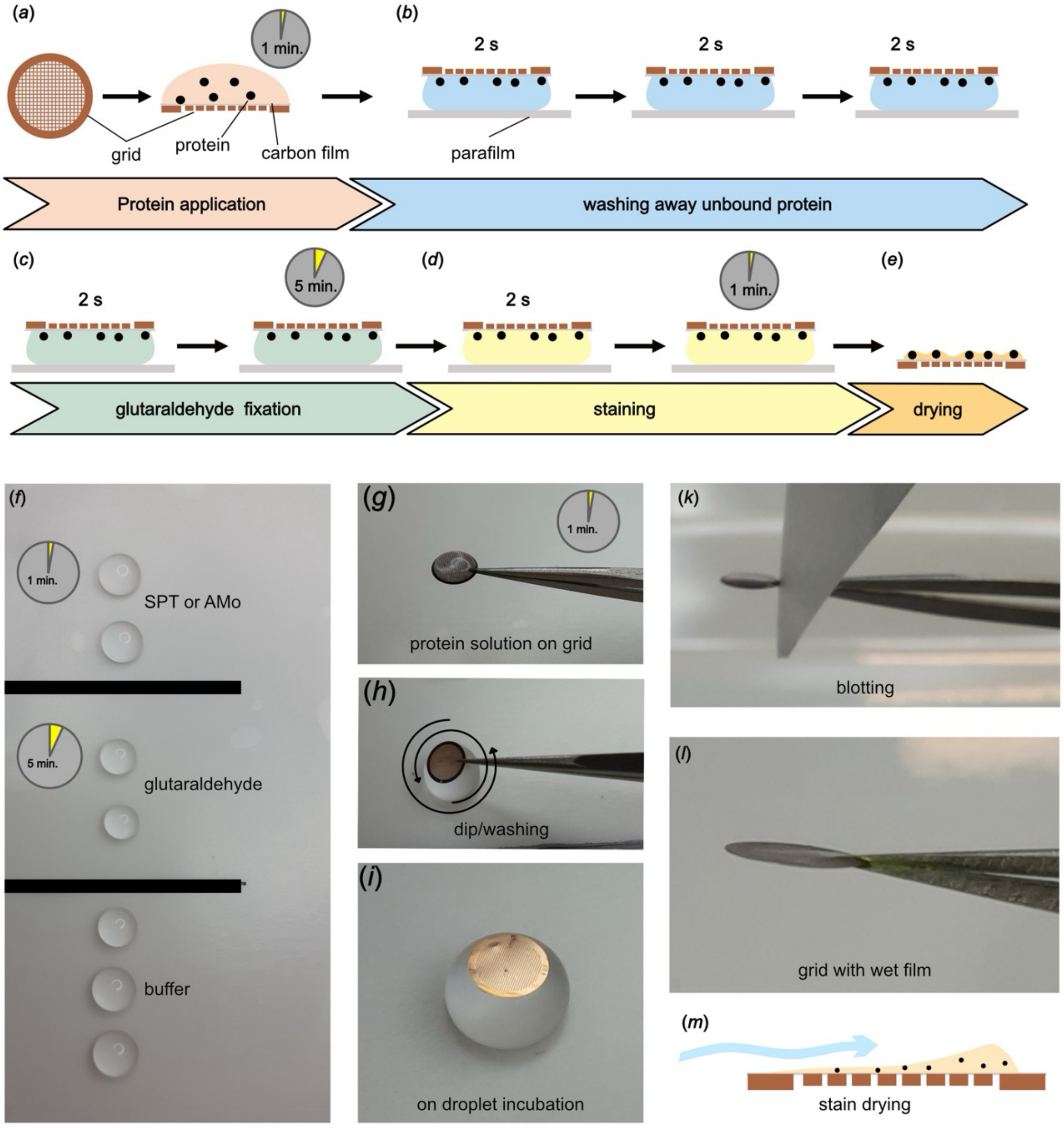


**Figure S2** **Detailed negative staining workflow**. (*a-d*) Cartoon workflow summary. (*a*) The protein solution is applied to a glow discharged sample carrier grid with a continuous support film and incubated for 1 min. (*b*) Grids are then washed with sample buffer. (*c*) Next, an on-grid fixation step using 0.15% glutaraldehyde solution is employed to fix the protein. (*d*) The protein is stained with either SPT or AMo. (*e*) Afterwards the stain is rapidly dried to create an amorphous film around the sample. (*f-m*) Photographs of important steps. (*f*) droplets of buffer, glutaraldehyde and staining solution are placed on flat parafilm. (*g*) protein solution is incubated for 1 min on the grid, held by forceps. (*h*) The grid is shortly dipped or washed on the droplet. (*i*) A grid is incubated on a droplet of fixation solution. (*k*) The liquid is carefully blotted from the grid with filter paper. (l) A thin film of liquid is left on the grid. (*m*) The grid is dried by a gentle blow of a hair dryer to create a gradient from thin to thicker stain on the grid.

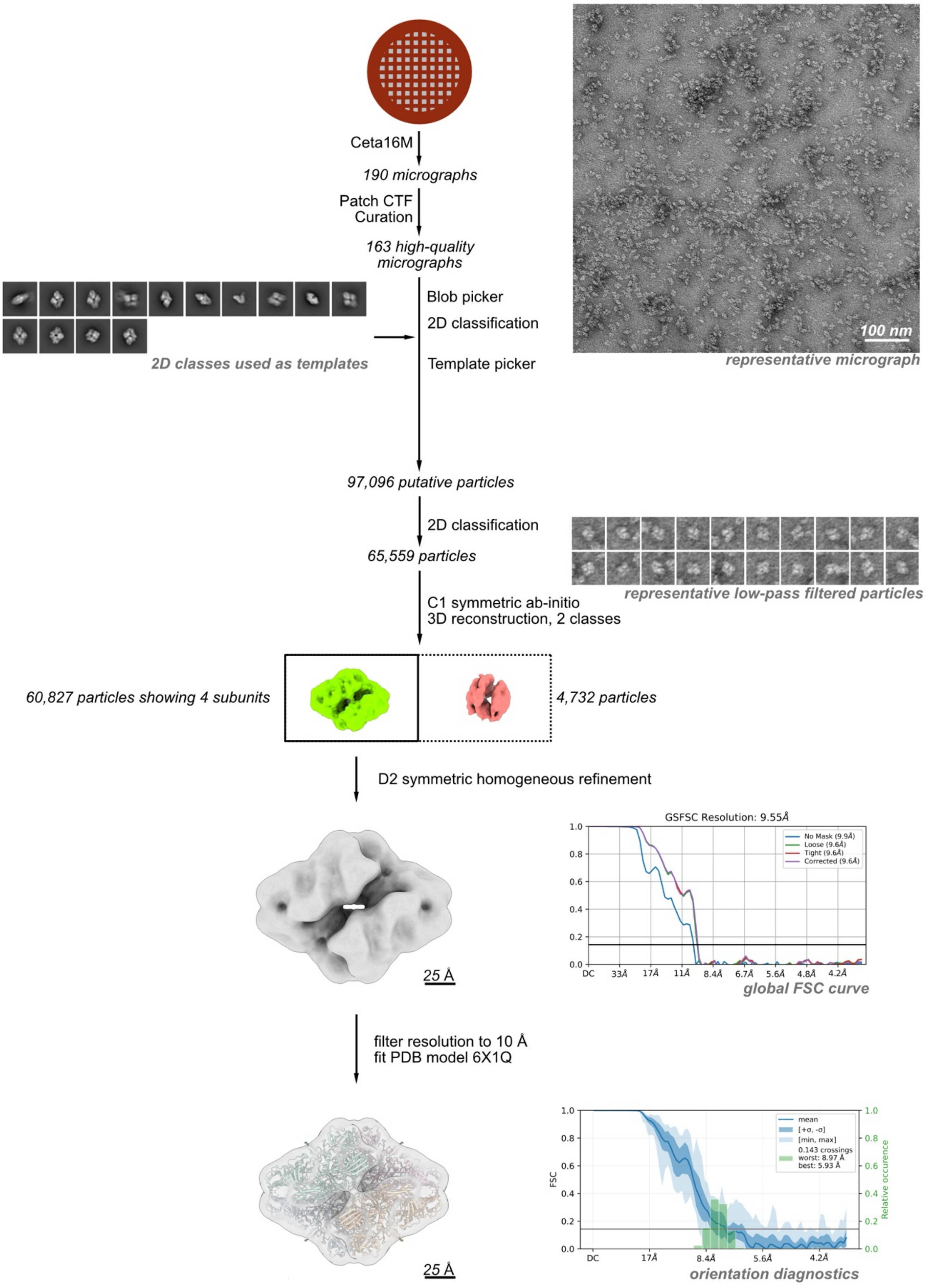


**Figure S3** **Reconstruction workflow for UF stained β-galactosidase.** A total of 190 micrographs were collected on a 120 kV electron microscope. From these 163 high-quality micrographs were selected based on good CTF scores and low astigmatism. Particles were first picked using a blob picker, and then subjected to unsupervised 2D classification. Representative classes showing protein-like structures were used for a template picker. Detected putative particles were curated using unsupervised 2D classification, selecting for particles with protein-like density. The selected particles were further curated using C1 symmetric ab-initio reconstruction, sorting them into 2 distinct populations. From these, all particles contributing to a structure showing clear density for 4 distinct subunits (shown in green and highlighted by a thicker box outline) were combined and refined in 3D using a homogeneous refinement algorithm enforcing D2 symmetry, resulting in a map with a uniform resolution of 9.6 Å. Before fitting of the atomic model, the map was low-pass filtered to 10 Å.

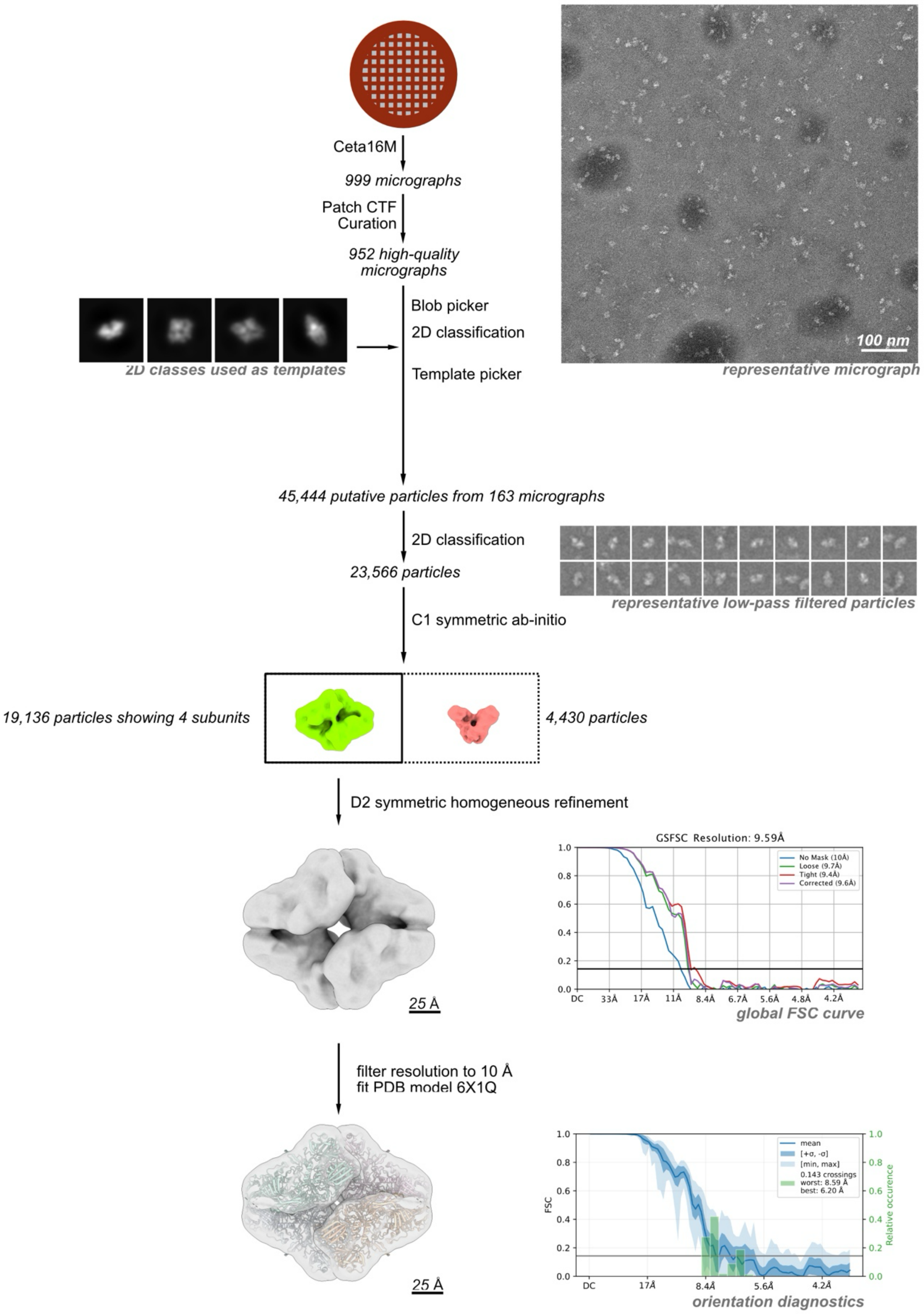


**Figure S4** **Reconstruction workflow for SPT stained β-galactosidase.** A total of 999 micrographs were collected on a 120 kV electron microscope. From these 952 high-quality micrographs were selected based on good CTF scores and low astigmatism. Particles were first picked using a blob picker, and then subjected to unsupervised 2D classification. Representative classes showing protein-like structures were used for a template picker. Detected putative particles on a random subset of 163 micrographs were curated using unsupervised 2D classification, selecting for particles with protein-like density. The selected particles were further curated using C1 symmetric ab-initio reconstruction, sorting them into 2 distinct populations. From these, all particles contributing to a structure showing clear density for 4 distinct subunits (shown in green and highlighted by a thicker box outline) were combined and refined in 3D using a homogeneous refinement algorithm enforcing D2 symmetry, resulting in a map with a uniform resolution of 9.6 Å. Before fitting of the atomic model, the map was low-pass filtered to 10 Å.

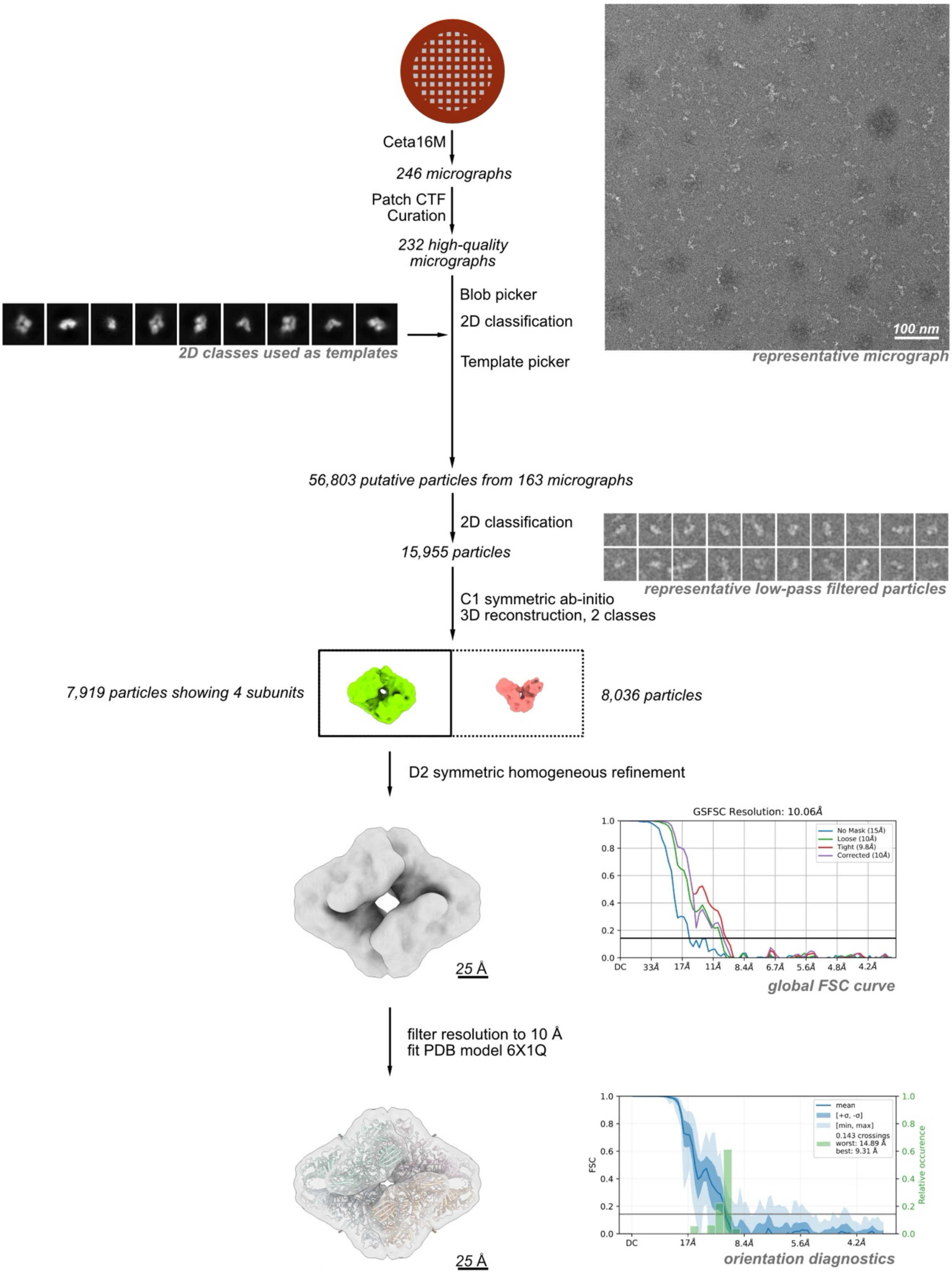


**Figure S5** **Reconstruction workflow for AMo stained β-galactosidase.** A total of 246 micrographs were collected on a 120 kV electron microscope. From these 232 high-quality micrographs were selected based on good CTF scores and low astigmatism. Particles were first picked using a blob picker, and then subjected to unsupervised 2D classification. Representative classes showing protein-like structures were used for a template picker. Detected putative particles on a random subset of 163 micrographs were curated using unsupervised 2D classification, selecting for particles with protein-like density. The selected particles were further curated using C1 symmetric ab-initio reconstruction, sorting them into 2 distinct populations. From these, all particles contributing to a structure showing clear density for 4 distinct subunits (shown in green and highlighted by a thicker box outline) were combined and refined in 3D using a homogeneous refinement algorithm enforcing D2 symmetry, resulting in a map with a uniform resolution of 10.6 Å. Before fitting of the atomic model, the map was low-pass filtered to 10 Å.

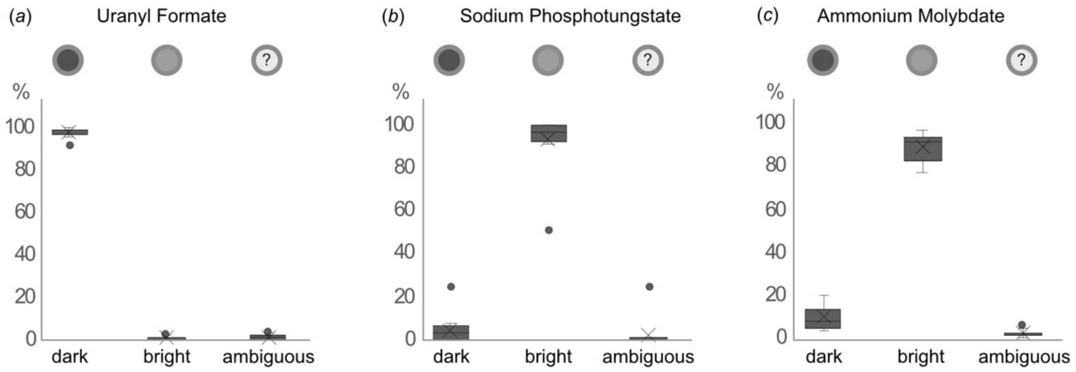


**Figure S6** **SPT and AMo differ in their staining behavior of apoferritin compared to UF.** Box plot diagrams of apoferritin stained with (*a*) UF, (*b*) SPT, (c) or AMo. For each staining conditions 5 images from 3 independent staining experiments were analyzed and observed apoferritin particles classified as either having a dark or a bright core. If unclear, particles were classified as ambiguous. The standard deviation was calculated from the differences between the three independent repeats. For UF, 98 ± 2 % of the particles were classified as dark, 0.7 ± 1 % as bright, with 1.3 ± 1.2 % ambiguous particles. For SPT, 4.3 ± 6.3 % of the particles were classified as dark, 93,8 ± 12.3 % as bright, with 1.9 ± 6.3 % ambiguous particles. For AMo, 9.4 ± 5.1 % of the particles were classified as dark, 89 ± 6.3 % as bright, with 1.6 ± 1.7 % ambiguous particles.

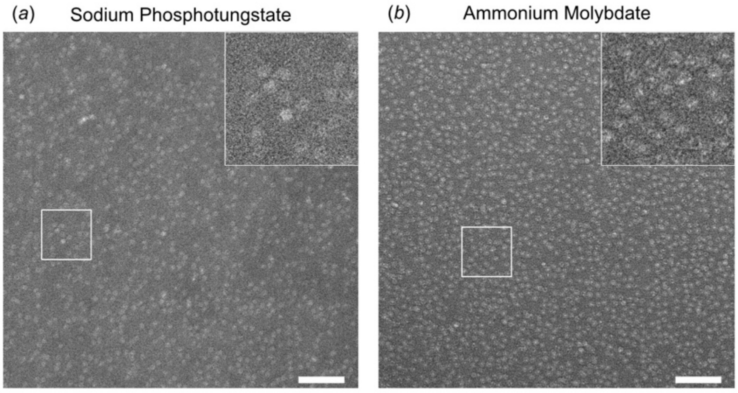


**Figure S7 Without a fixation step, SPT and AMo stained apoferritin particles have poor contrast.** Representative raw micrographs of apoferritin stained with (*a*) SPT, (*b*) or AMo. Scale bars are 100 nm. The insets show the boxed area at 2.5 x magnification. For micrographs of SPT and AMo staining with an additional on-grid fixation see Figure 1.
